# Refinement of the primate corticospinal pathway during prenatal development

**DOI:** 10.1101/575936

**Authors:** Ana Rita Ribeiro Gomes, Etienne Olivier, Herbert P. Killackey, Pascale Giroud, Michel Berland, Kenneth Knoblauch, Colette Dehay, Henry Kennedy

## Abstract

Perturbation of the developmental refinement of the corticospinal pathway leads to motor disorders. In non-primates developmental refinement is well documented, however in primates invasive investigations of the developing corticospinal pathway have been confined to neonatal and postnatal stages when refinement is relatively modest.

Here, we investigated the developmental changes in the distribution of corticospinal projection neurons in cynomolgus monkey. Injections of retrograde tracer at the cervical levels of the spinal cord at embryonic day (E) 95 and E105 show that (i) areal distribution of back-labeled neurons is more extensive than in the neonate and dense labeling is found in prefrontal, limbic, temporal and occipital cortex; (ii) distributions of contra- and ipsilateral projecting corticospinal neurons are comparable in terms of location and numbers of labeled neurons, in contrast to the adult where the contralateral projection is an order of magnitude higher than the ipsilateral projection. Findings from one largely restricted injection suggest a hitherto unsuspected early innervation of the gray matter.

In the fetus there was in addition dense labeling in the central nucleus of the amygdala, the hypothalamus, the subthalamic nucleus and the adjacent region of the zona incerta, subcortical structures with only minor projections in the adult control.

## Introduction

The brain exerts control over movement of the body through descending telencephalic, hypothalamic, diencephalic, midbrain and hindbrain pathways (Kuypers 1982; Nudo and Masterton 1988; Lemon 2008). Among these parallel-descending routes, the corticospinal (CS) pathway provides the fastest and most direct influence of the cerebral cortex over different aspects of planning, execution and control of voluntary movements (Dum and Strick 1996, 2002; Lemon 2008).

Certain features of the adult and developing corticospinal system are conserved across species. In the adult the CS tract originates in layer 5 of the somatomotor regions of the frontal and parietal lobes. The axons of the large CS projection neurons descend ipsilaterally through the internal capsule, and in the lower brainstem most of the CS fibers decussate in the medullary pyramids, invading the contralateral spinal cord. Descending in the spinal funiculi, the CS fibers invade the spinal gray matter, establishing functional synapses with the interneurons, and, in some primates, motoneurons, located in different antero-posterior segments of the spinal cord. During development there is evidence of common maturational events involving large-scale elimination of CS projections (Stanfield 1992; Luo and O’Leary 2005; Martin 2005). Firstly, there is evidence of a refinement of the spatial extent of CS fiber innervation of spinal gray matter as development progresses (hamster: Reh and Kalil (1981); cat: Alisky et al. (1992), Li and Martin (2000); opossum: Cabana and Martin (1985); macaque: Kuypers (1962), Armand et al. (1997)). Secondly, there is a widespread distribution of CS projection neurons in the cortex at early stages of development and a subsequent elimination of CS axons leading to the substantially more restricted pattern of CS projection neurons observed in the adult (mouse: Kamiyama et al. (2015), rat: Stanfield et al. (1982), Leong (1983), Bates and Killackey (1984), Schreyer and Jones (1988), ferret: Meissirel et al. (1993), opossum: Cabana and Martin (1984), macaque: Galea and Darian-Smith (1995)).

Across species, the adult CS pathway exhibit important structural differences, particularly between primates and non-primates. In primates there is a species-specific increase in: (i) the number and size of the CS fibers; (ii) the percentage of fibers projecting to the ipsilateral spinal cord, either by not crossing in the pyramidal decussation or by re-crossing the midline in the spinal cord; (iii) importantly, the primate CS pathway is characterized by monosynaptic projections from the cortex to the motoneurons in the gray matter of the spinal cord (Lemon 2008; Welniarz et al. 2017). This well-developed feature in Old World monkeys, great apes and man is thought to be essential for the acquisition of specialized motor functions. In particular, cortico-motoneuronal connections are key for the development of unique levels of manual dexterity including the capacity to achieve varying degrees of independent finger movements (IFM) (Lemon 2008).

Regarding the developmental refinement of the CS pathway, previous studies in primates mostly addressed the maturation of axonal terminations, correlating the development of fine motor skills such as IFM, with the postnatal increase of motoneuron pool innervation by the cortex (Lawrence and Hopkins 1976; Armand *et al.* 1997; Olivier et al. 1997). There is a paucity of studies using invasive techniques addressing the early widespread distribution of cortical neurons projecting to the spinal cord. To our knowledge, there is only one study specifically addressing this issue in primates, and specifically in the macaque. By examining the topographical changes in the distribution CS projection neurons during the first postnatal year, Galea and Darian-Smith (1995) showed a threefold reduction in the number of CS projection neurons and a 50% reduction of the cortical territory where these cells were located. While these findings confirm the presence of a wider projection prior to adult maturation, the results obtained differed significantly from those obtained in non-primates. Whereas in the neonate macaque the early projections (ipsi- and contralateral) are overall restricted to the same frontal, parietal and insular regions that project to the spinal cord in the adult, tract tracing studies in the rat show that during development there are cortical neurons throughout the neocortex projecting to the spinal cord regions (Stanfield *et al.* 1982; Leong 1983; Bates and Killackey 1984; Schreyer and Jones 1988; Joosten and van Eden 1989).

These results raise the question of whether the relatively spatially confined pattern observed perinatally in the macaque (i) reflects a primate-specific feature, suggesting that in primates the initial transient projection is considerably more restricted than the projection observed during early development in other species; or alternatively (ii) if extensive transient projections do exist, but are largely eliminated before birth, and hence not observed in the postnatal period.

The development of other cortical projection systems in primate could seem to support the idea of an early patterned connectivity in this order. The complete absence during primate development of widespread callosal connections between the primary visual cortices (Dehay et al. 1988; Chalupa et al. 1989) suggests the possibility of a high degree of connectional specificity during primate development. Just how generalized is this connectional specificity is an open question, particularly pertinent to the CS system in which a wealth of perinatal clinical issues could implicate complications in developmental refinement of the CS pathway. For instance, there is clinical evidence suggesting the presence of exuberant CS projections; trans-magnetic stimulation (TMS) of the occipital cortex of an infant at 14 and 48 months who had a prenatal stroke, elicited motor movements with the same characteristics as those elicited by stimulation of sensorimotor cortex (Basu et al. 2010). These observations suggest either that the infant suffered from an anomalous projection or that it had not undergone the normal developmental refinement of a transient pathway.

In the present study we have investigated the development of the CS pathway in the prenatal macaque. Use of retrograde tracer injections in the cervical spinal cord revealed a widespread distribution of CS projection neurons from both contra- and ipsilateral hemispheres. In one instance the injection site was largely restricted to the spinal gray matter and appeared to spare the descending fiber pathways in the spinal cord. The results from this injection suggest that invasion of the gray matter could be more extensive than previously suspected. In addition we found extensive labeling of subcortical structures in the fetus that is much restricted during later development.

## Material and Methods

The present study is based on observations in two fetuses and one adult control (**Table 1**). Surgical and histology procedures were in accordance with European requirements 86/609/EEC.

**Table 1:** Experimental cases. E embryonic day, FB Fast Blue.

| **Case** | **Age at injection** | **Age at perfusion** | **Injection site** | **Tracer** | **Tracer Volume** | **Plane of section** |
| --- | --- | --- | --- | --- | --- | --- |
| BB152 | E95 | E108 | C1 | FB | 0.6 μl | Parasagittal |
| BB171 | E105 | E117 | C1 | FB | 0.3 μl | Parasagittal |
| Adult | 10 years | 10 years | C4 | FB | 4 μl | Parasagittal |

### Injections of retrograde tracers in the fetus

Timed pregnant cynomolgus monkeys (Macaca fascicularis) received atropine (1.25 mg, i.m.), dexamethasone (4 mg, i.m.), isoxsuprine (2.5 mg, i.m.), and chlorpromazine (2 mg/kg, i.m.) surgical premedication. They were prepared for surgery under ketamine hydrochloride (20 mg/kg, i.m) anesthesia. Following intubation, anesthesia was continued with 1-2% halothane in a N20/02 mixture (70/30). The heart rate was monitored, and the expired CO2 maintained between 4.5% and 6%. Body temperature was maintained using a thermostatically controlled heating blanket. On embryonic day 95 (E95) and E105, and using sterile procedures, a midline abdominal incision was made, and uterotomy was performed. The fetal head and neck was exposed, and the tracer Fast Blue (FB, 0.3–0.6 μl of aqueous solutions at 3%) injected in the lateral cervical spinal cord under visual inspection with a Zeiss binocular surgical microscope control and using glass micropipettes. The fetus was replaced in the uterus and the incisions were closed. The mother was returned to her cage and given an analgesic (visceralgine, 1.25 mg, i.m.) twice daily for 2 days.

The pregnant monkey received postoperative medication consisting of a muscular relaxant (ixosuprine chlorhydrate) and an analgesic (tiemonium methylsulfate). Fetuses were delivered by caesarian section after a 12-13 day survival period. Animals were deeply anaesthetized before being perfused transcardially with 200 ml of 2.7% saline, 1-3 liters of a 0.5% glutaraldehyde and 8% paraformaldehyde mixture in 0.1 M phosphate buffer, 0.5 liters of 8% sucrose, 0.5 liter of 20% sucrose, and 0.5 liter of 30% sucrose in phosphate buffer.

### Injections of Retrograde Tracers in the adult control

Identical medication, anesthesia and monitoring procedures were used as described above. Tracer injections were placed using a Zeiss binocular surgical microscope. Injections were made by means of Hamilton syringes in a stereotypic fashion. Following injections, artificial dura mater was applied, the bone flaps closed, cemented and the scalp stitched back into position.

Fourteen days after injections to allows optimal retrograde labeling of neurons projecting to the pick-up zone, the adult animal was anesthetized with ketamine (20 mg/kg, i.m.) followed by a lethal dose of Nembutal (60 mg/kg, i.p.) and perfused through the heart with a 8% paraformaldehyde and 0.05% glutaraldehyde solution. After fixation, perfusion was continued with 10, 20 and 30% sucrose solutions to provide cryoprotection of the brain.

### Data Acquisition

Following cryoprotection the brains and spinal cords of the fetuses and the brain of the adult control were immediately removed and blocked. Forty micrometers thick sections cut on a freezing microtome and mounted in saline onto gelatinized slides. The brains were cut parasagitally and the spinal cords horizontally. One in 3 sections of the adult brain and 1 in 4 sections in the fetuses were retained to explore CS neuron distribution. Sections were examined in UV light with oil-immersion objectives using a Leitz fluorescence microscope equipped with a D-filter set (355-425 nm). High precision maps were made using Mercator software running on Exploranova technology, coupled to the microscope stage. At least one in every ten sections (1.6mm apart in the fetuses and 1.2mm in the adult) was observed and labeled neurons charted across the cortex and subcortical structures. The frequency of charting was increased to 1 in every 2 section in order to have a complete sampling of layer 5 and subcortical midline structures.

Controlled high frequency sampling gives stable neuron counts despite curvature of the cortex and heterogeneity of neuron distribution in the projection zones of individual areas (Vezoli et al. 2004; Markov, Vezoli, et al. 2014).

Selections of sections were processed for cytochrome oxidase (CO) and acetylcholine esterase (AChE) using published procedures (Hardy et al. 1976; Silverman and Tootell 1987). Characteristics of neurons labeled with Fast Blue are described by Keizer and colleagues (1983). Area and/or regional limits were reported on the charts of labeled neurons (**Table S1**). Whenever possible, these neurons were then attributed to areas of our atlas (**Table S1**) based on landmarks and histology, and counted according to that parcellation (Markov, Ercsey-Ravasz, et al. 2014).

### Quantification of projections and statistical analysis

All statistical analyses were performed in the R statistical environment (R Development Core Team 2016). We defined the strength or weight of a projection from a given region with respect to the number of labeled neurons in that region. Because the number of labeled cells depends on the amount of tracer injected in each spinal cord, referring to the total number of neurons of each projection is not appropriate. Hence, we calculated a normalized weight index for each projection using the Fraction of Labeled Neurons (FLN). As previously defined (Markov et al. 2011), the FLN is the proportion of cells located in a given source region with respect to the total number of labeled neurons in the cortex. At all ages, FLN values were computed using the interpolated number of neurons in each region, which is obtained by inferring the numbers of labeled neurons on fluorescence sections that were not examined.

## Experimental findings

Injections of tracer were made to one side of the spinal cord so that we could determine the frequency of labeled CS neurons separately for the ipsi- and contralateral hemispheres. Injections were made in two fetuses, the youngest was injected at E95 and perfused at E108, and the elder was injected at E105 and perfused at E117 (**Table 1**). In the description of the results we shall refer to these two cases by the age of perfusion. In a first instance we address the histological location of the injection sites. Next, in order to determine the areal location of CS projecting neurons in the fetuses we describe how we identified the areal borders with respect to the evolving sulci pattern. In order to obtain an accurate understanding of the changing areal pattern of CS neurons we have made counts of labeled neurons, which we first describe in the adult before proceeding to report our findings in the fetuses. Finally, we briefly describe labeled spinal cord projection neurons in the subcortical structures in fetuses, which are considerably more numerous than in adult counterpart, suggesting that they constitute bona fide exuberant projections.

### Location of the injection sites

Injections targeted the right side of the spinal cord near the midline in order to maximally involve the spinal gray matter. Injections of Fast Blue gave rise to a dense and restricted tracer deposit. An important characteristic of the fluorescent tracer Fast Blue is that we have a relatively precise understanding of its pick-up zone (i.e. the spatial extent of the region in which tracer is taken up by axons and transported back to cell bodies) (Bullier and Kennedy 1983; Bullier et al. 1984). The pick-up zone corresponds to a region of dense extracellular dye (**Figure 1B**). This made it possible to accurately determine the full extent of the pick-up zone in the E108 and E117 fetuses, which in both cases was immediately below the pyramidal decussation.

**Figure 1:**
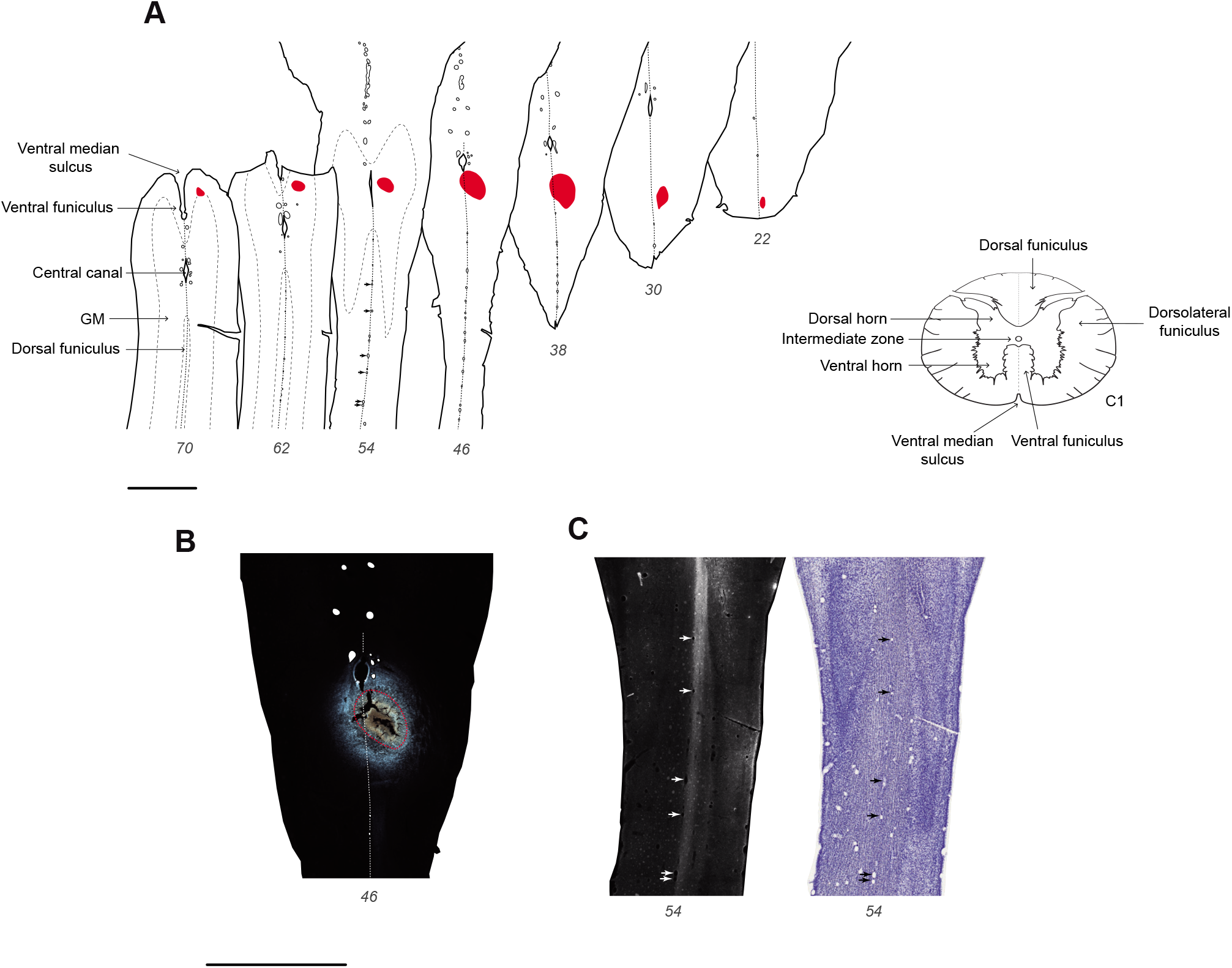
Fluorescent micrograph of E117 injection site. **A:** horizontal sections of the upper spinal cord with the pick-up zone of the Fast Blue injection site indicated in red; **B**: fluorescent micrograph showing the pick-up zone outlined in red in section 46 in panel A; **C**: high-exposure micrograph caudal to the injection site in section 54 in panel A, showing labeled fibers limited to the injected side of the spinal cord (see **Figure S1** for further detail). Black and white arrows highlight the blood vessels present on the midline of the spinal cord. Scale bar: 2mm

In the E117 fetus the pick-up zone involved the spinal intermediate and ventral gray matter and the dorsal funiculus (**Figure 1**). Importantly, this injection spared the white matter compartments known as the lateral and ventral dorsal funiculi where the descending fiber pathways are located. Hence, we hypothesize that the labeling from this injection results from pick-up and transport from axons that have invaded the gray matter of the spinal cord *(see Discussion).*

Reconstruction of the E117 injection site showed a minimal 150-micron intrusion of the pick-up zone in the gray matter of the left side of the cord (see panels 38 and 46 in **Figure 1A and B**). This Figure shows that there is a conspicuous labeling of the dorsal funiculus running posterior to the injection site, which was observed to be restricted to the right side of the spinal cord (**Figure 1C**). Importantly, we conclude that the minute involvement of the left cord was insignificant as it failed to label the equivalent longitudinal tract on the left side of the spinal cord (**Figure 1C and S1**). These observations suggest that the pick-up zone in this injection was very largely restricted to the right hand gray matter of the spinal cord.

Inspection of the injection site in the E108 fetus in **Figure 2** reveals an extended pickup zone in the right pyramidal tract involving the gray matter. However, there was also an unexpected, limited, secondary pick-up zone restricted to the gray matter-white matter boundary of the left side of the spinal cord (**Figure 2**). In the adult, FB was injected at the level of cervical vertebrae C4. In this case, no histology of the injection site in the spinal cord was available, and its location was inferred from the distribution of labeled CS neurons (see *Discussion*).

**Figure 2:**
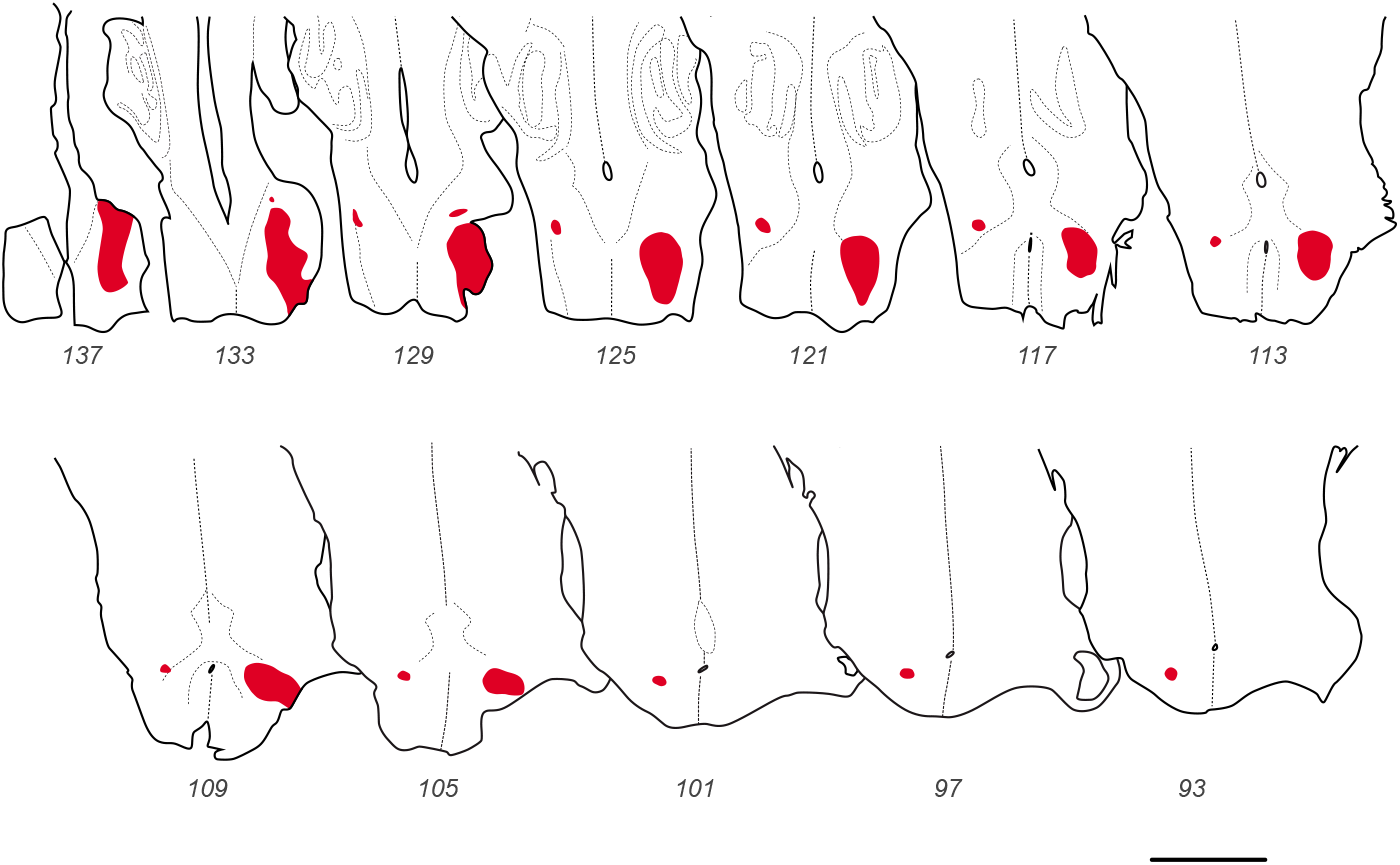
E108 Injection site. Horizontal sections of the upper spinal cord and medulla with the pick-up zone of the injection site indicated in red. Scale bar: 2 mm.

### Regional and presumptive areal boundaries with respect to the developing sulci pattern

In order to determine the developmental changes of CS neurons in the two fetuses and compare this with the adult distribution, it was necessary to parcellate the cerebral cortex across ages into comparable regions with respect to stable landmarks. In the fetal macaque there is a gradual transition from a lissencephalic to a gyrencephalic brain (Sawada et al. 2012). Hence, the use of cortical gyri and sulci as landmarks for areal and regional parcellation is relatively limited particularly at early developmental stages.

At E108 there are five clearly identifiable sulci (**Figures 3A and B**, and left panels in **Figures 4 and 5**): the lateral (ls), the parieto-occipital (pos), the superior temporal (sts), the circular and the hippocampal sulci. Previous studies have reported the earliest manifestations of these sulci being between E70 to E100 (Sawada *et al.* 2012). In addition, we were able to detect the presumptive calcarine (cas), central (cs), arcuate (ars), inferior occipital (ios), occipitotemporal (ots), and anterior middle temporal sulci (amts), which have been reported to appear respectively at E80, E90, E90, E90, E120 and E120 (Sawada *et al.* 2012).

**Figure 3:**
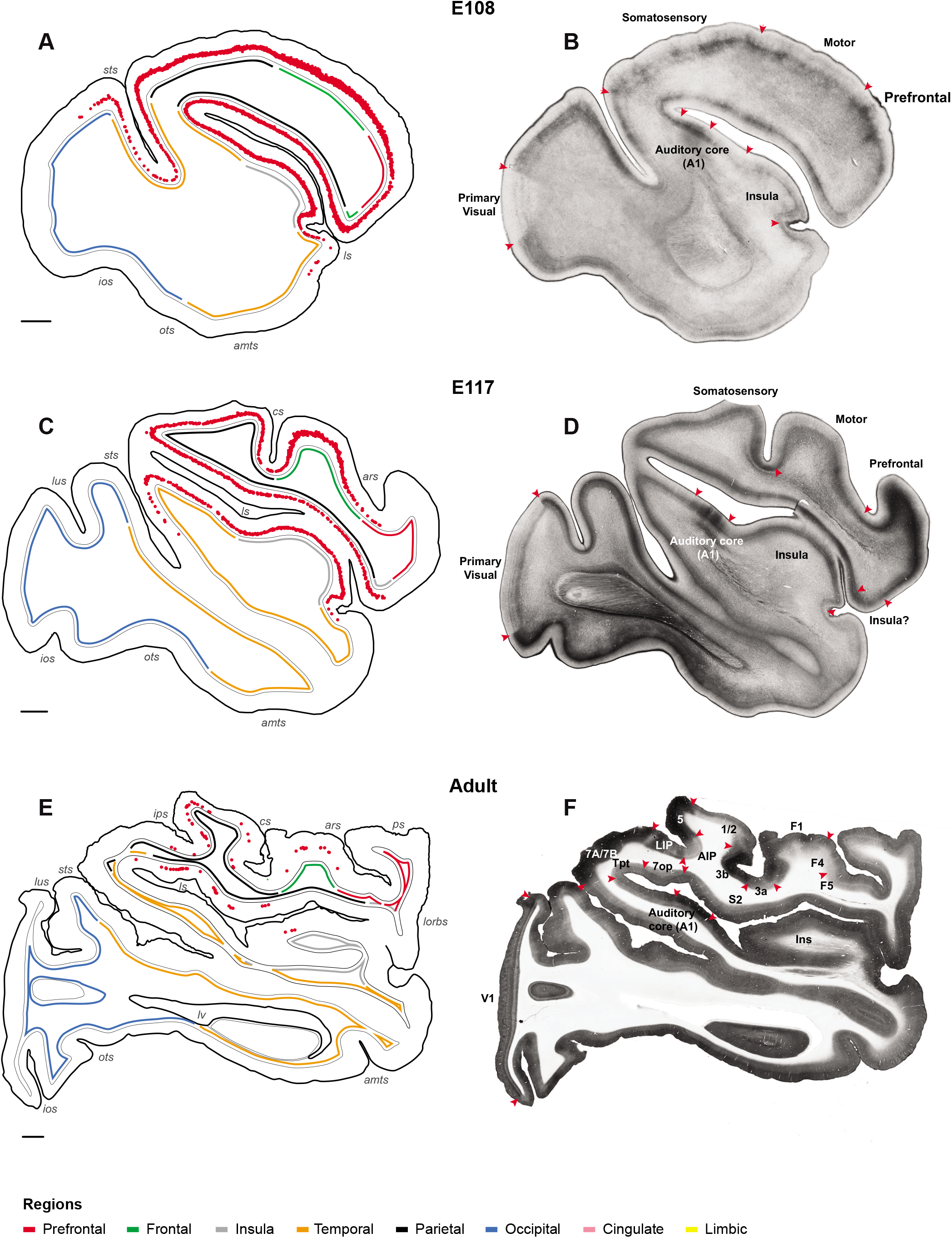
Regional boundaries at E108, E117 and in the adult. **A, C** and **E**: Distribution of retrograde labeled neurons (red dots) in the left hemisphere in parasagittal sections at equivalent lateral-medial levels following injection in the right cervical spinal cord. Colored lines indicate the different regions (red, prefrontal; green, frontal; gray, insula; orange, temporal; black, parietal; blue, occipital; pink, cingulate; yellow, limbic). **B, D** and **F**: presumptive regional and areal boundaries identified with AChE histochemistry in E108 and E117 and CO histochemistry in the adult. Note the well-defined boundaries of primary visual (V1) and auditory (Core) cortex. For areal abbreviations in this and subsequent Figures, see **Table S1**. Scale bar: 2 mm.

The older E117 fetus exhibited a significantly more mature sulci pattern, as nearly all the sulci found in the adult could be identified at this stage (**Figures 3C and D**, and middle panels in **Figures 4 and 5**). In addition to those observed in the youngest fetus, at E117 we could identify the lunate (lus), the intraparietal (ips), and the principal sulci (ps), which have been reported to appear between E100 and E110 (Sawada *et al.* 2012). Sulci that were only very weakly formed at E108, such as the central and arcuate sulci, were much more marked in the older fetus.

**Figure 4:**
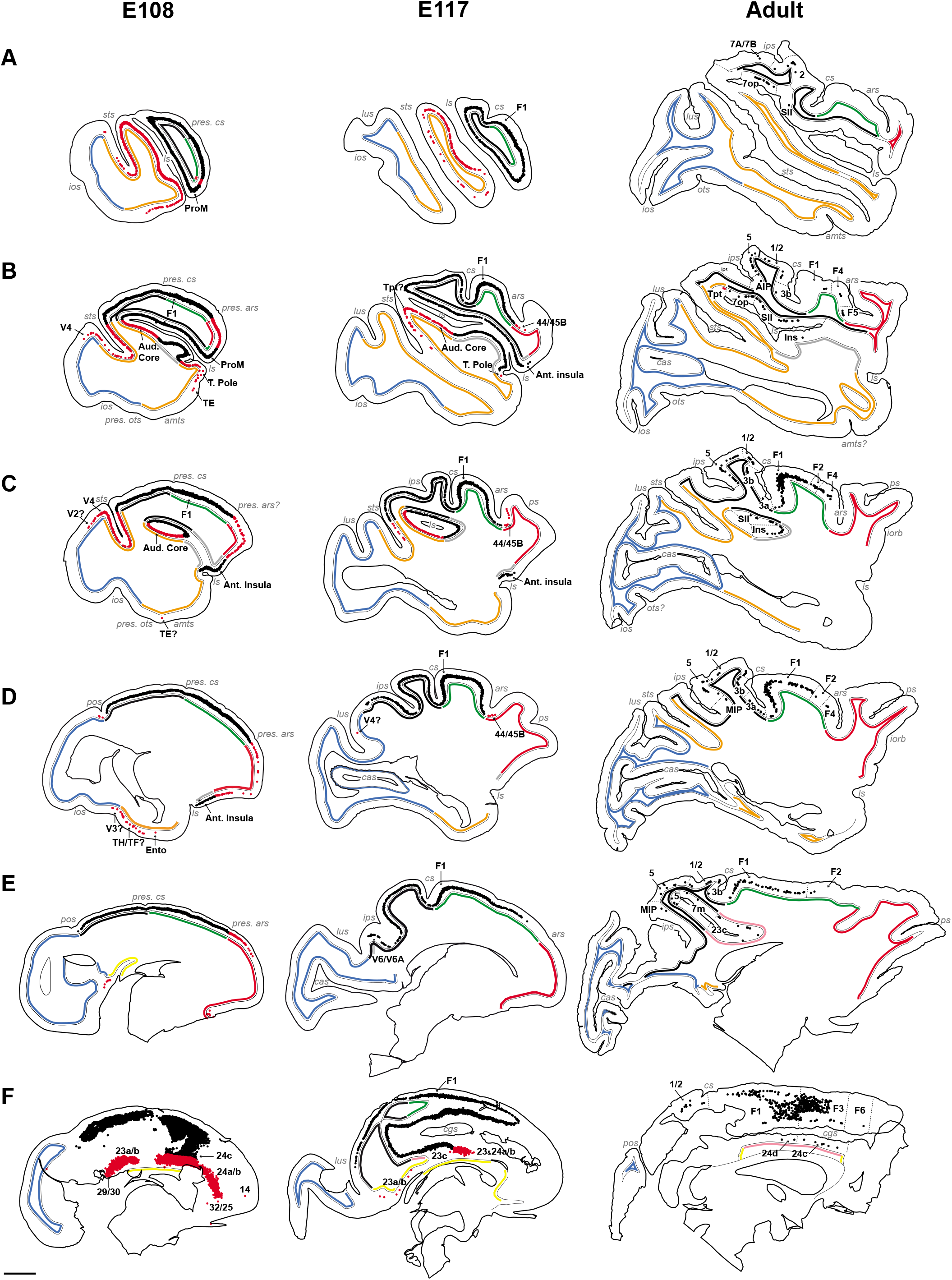
Developmental refinement of contralateral CS projecting neurons. Sections of left hemisphere at equivalent medio-lateral levels across all three ages showing the areal distribution of CS pathway projection neurons. Labeled neurons in cortical regions that maintain CS neurons in the adult are in black, labeled neurons in regions that lose their projection to the spinal cord in red. Colored lines indicate different region (see color code in **Figure 3**). Scale bar: 5 mm.

### Cortical landmarks and cytoarchitecture

**Figure 3** shows exemplar sections displaying the distribution of CS neurons contralateral to the injection site (i.e. in the left cortical hemisphere) following injection in the right cervical cord in the fetuses as well as in the adult control. Individual red dots represent FB back-labeled cells. Here and in following figures, the cortex in all three brains was subdivided into eight regions. Colored lines beneath the gray matter-white matter border code the regional location of labeled neurons: prefrontal (red), frontal (green), insular (gray), temporal (orange), parietal (black), occipital (blue), cingulate (pink) and limbic (yellow) (see **Table S1** for individual area allocation).

The adult brain was parcellated according to a 91-area atlas that combines histological criteria with defined landmarks (Markov, Ercsey-Ravasz, *et al.* 2014). Because the present study uses a parasagittal plane of section, we converted the atlas from the coronal plane into the parasagittal plane using a 7T MRI scan of the contralateral hemisphere used to create the Markov, Ercsey-Ravasz *et al*. (2014) atlas. ITK-SNAP software (http://www.itksnap.org) made it possible to create the virtual parasagittal plane displaying the appropriate area locations. Histological boundaries were confirmed using Nissl, CO and AChE stained sections.

Boundary locations in the developing brains were checked using Nissl (not shown) and AChE stained sections (**Figure 3**). In the youngest fetus the primary visual cortex is readily apparent as a region of very low levels of AChE activity. At more anterior levels presumptive auditory cortex is relatively strongly labeled by AChE. In the E117 fetus, the transition from parietal to frontal cortex in the central sulcus coincided with a decrease in the density and thickness of layer 4 and AChE activity in the frontal cortex. The transition from frontal to prefrontal cortex in the presumptive arcuate sulcus was accompanied by an increase in AChE expression in the prefrontal cortex. The transition from temporal to occipital cortex was indicated by an increase in the thickness of layer 4 and an increase of AChE activity in the occipital cortex. The anterior insular cortex in the posterior orbitofrontal cortex was identified by a decrease in layer 4 thickness and lower AChE activity when compared to neighboring regions. The remaining boundaries were determined using sulcal landmarks.

### Contralateral projecting neurons

At all ages, the FB back-labeled cells in the cortex are layer 5 pyramidal neurons. In the adult, CS neurons projecting contralaterally are located principally in the frontal lobe and to a lesser extent in the parietal and in the cingulate regions (**Figure 6**, see **Figure 4**, right column). In the frontal cortex CS neurons are found in the primary motor cortex (area F1), and in all six subdivisions of the premotor region within the lateral dorsal (areas F2, F7), lateral ventral (areas F4, F5), and mesial (areas F3, F6) cortex where they are restricted to the arm, leg and trunk representations (Cure and Rasmussen 1954). The maximum density of CS neurons is on the lip of central sulcus containing area F1 and the representation of movements of the fingers (**Figure 4, rows C and D**). In parietal cortex CS neurons are observed in areas 3a, 3b, 1/2, 5, 7A/7B, AIP, MIP, LIP, 7op and SII (rows A to E).

The peak levels of CS neurons in the primary somatosensory area are reported to be located within the representation of the forelimb (wrist and arm) (Kaas et al. 1979). CS neurons are also observed in the insular cortex (rows B and C) throughout its granular subdivision, principally in the circular sulcus containing the representation of body parts (Robinson and Burton 1980). In addition, CS neurons are observed in multiple limb representations in the cingulate cortex in areas 23c, 24c and 24d (He et al. 1995) (**rows E** and **F**).

During development we observed a massive reduction in the cortical territory containing labeled CS neurons (**Figure 4**). In the youngest fetus, outside of the occipital and frontal poles, labeled CS neurons form a near continuous territory spanning many regions lacking CS neurons in the adult. The older fetus showed a small reduction in the spatial extent of CS neurons, which, nevertheless, continue to have a considerably wider distribution compared to that observed in the adult.

**Figure 4** distinguishes the distribution of early-formed CS neurons, which according to their location, are destined to be lost in development (marked in red), from those CS neurons that are located in regions where CS neuron are found in the adult (marked in black). This figure reveals the changing distribution of CS neurons projecting to the contralateral spinal cord during development. Those regions which in the adult will contain CS neurons (i.e. black dots in **Figure 4**) exhibit a continuous band of CS neurons during fetal development in contrast to the discontinuous distribution observed in the adult. The regions that project transiently (i.e. red dots) during development, namely temporal (orange line), occipital (blue line), prefrontal (red line) and limbic (yellow line) regions exhibit extensive labeling.

Whereas the adult temporal cortex virtually sends no projections to the spinal cord (excepting 2 cells in Tpt; **Figure 4 row B**), transient CS projections from the temporal cortex were numerous during development. At E108 and E117, CS neurons were observed in many subdivisions of the temporal cortex including the posterior bank of the lateral sulcus where the primary auditory cortex (area A1) and belt auditory cortex is located in the adult (**rows A** to **C**). In addition, CS neurons in the fetuses were observed in the anterior bank of superior temporal sulcus, within the polysensory area STP. In the youngest fetus, transient CS neurons in the temporal cortex were more extensive and included (i) the temporal pole (**row B**), (ii) the fundus and posterior bank of the superior temporal sulcus, where presumptive areas TEa/m, TEOm, IPa, FST, MT and V4t are housed (rows A to C); (iii) the inferior temporal cortex, presumably within areas TE and TH/TF (rows A, B and D); and finally (iv) in the subiculum, sparse labeling was observed (not shown). Overall contralateral temporal cortex contributes 11.7% of the labeled CS neurons at E108 and 5.8% at E117.

The occipital cortex is devoid of CS neurons in the adult and only very sparse projections are present at E108 and virtually none at E117 (**Figure 4**). At E108, occipital CS neurons are present in presumptive areas V4 and V2 areas (**rows B** and **C**) and a few labeled neurons are observed in presumptive contralateral V3 (row D). At E117, occasional labeled CS neurons are located in presumptive V4 (**row D**). Note that at E108 and E117 no back-labeled CS neurons were found in the primary visual area, area V1. Overall occipital cortex contributes 0.3% of labeled CS neurons at E108 and 0.02% at E117.

In the prefrontal cortex there are numerous CS neurons during development that disappear entirely in the adult brain. Between E108 and E117 there is a reduction in the extent of CS neurons (**Figure 4, rows D** and **E**). At E117, only a sparse projection is observed immediately anterior to the frontal-prefrontal boundary in the arcuate sulcus (rows B to D), in presumptive areas 44, 45B, and FEF. Overall prefrontal cortex contributes 2.3% of labeled CS neurons at E108 and 0.33% at E117.

Limbic CS neurons arise from presumptive areas 23a/b, 24a/b and 32/25 within the cingulate gyrus, in both E108 and E117 (**Figure 4, rows F**). Additional CS neurons are observed in presumptive retrosplenial cortex, areas 29/30, at E108 (**row F**). Overall limbic cortex contributes 7.03% of the labeled CS neurons at E108, and 1.14% at E117.

### Ipsilateral projecting neurons

**Figure 5** shows the distribution of ipsilateral CS neurons at approximately the same levels as those in **Figure 4**. Adult ipsilateral CS neurons are located in those cortical areas containing contralateral labeled neurons, but in the case of the ipsilateral projections labeled CS neurons are found in a more restricted territory (compare row D in **Figures 4** and **5**). As for the contralateral hemisphere, the main source of adult ipsilateral CS neurons is the frontal cortex. Adult parietal cortex has only sparse CS neurons, only slightly more numerous at medial levels (**Figure 5, row D**) and very few CS neurons in the insula (not shown) and cingulate cortices (**row E** and **F**).

**Figure 5:**
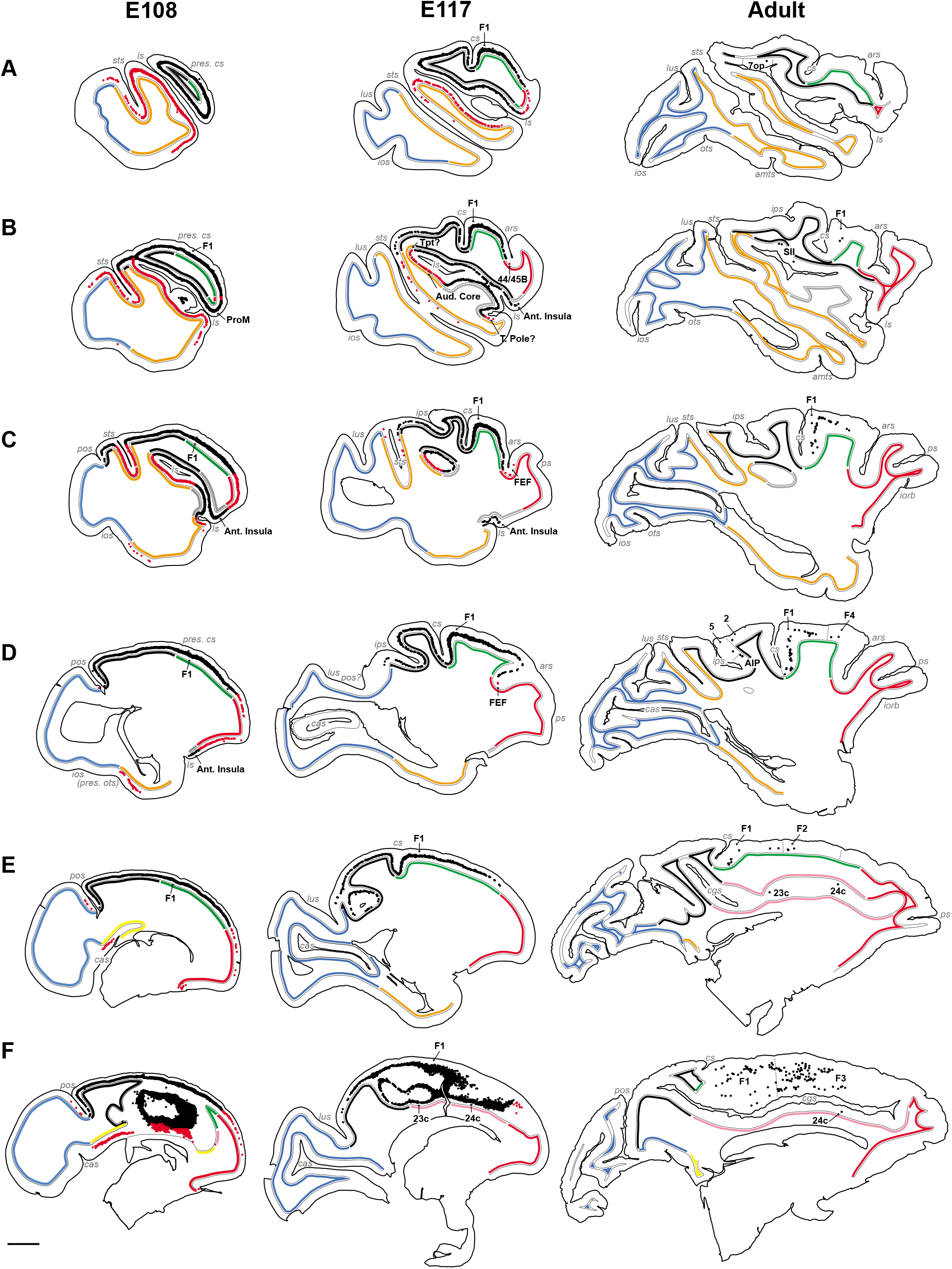
Developmental refinement of ipsilateral CS projecting neurons. Sections of the right hemisphere at equivalent medio-lateral levels across all three ages showing the areal distribution of CS pathway projection neurons. Labeled neurons in cortical regions that maintain CS neurons in the adult are in black, labeled neurons in regions that lose their projection to the spinal cord in red. Colored lines indicate different regions (see colored code in **Figure 3**). Scale bar: 5 mm.

Contrary to the adult, there is only a minimal lateralization of the CS tract during early development, so that the numbers and distribution of contra- and ipsilateral corticospinal neurons are much more comparable in the fetus compared to that observed in the adult. Whereas in the adult 20.1% of the back-labeled neurons were labeled in the ipsilateral hemisphere, in the E108 and in the E117 the ipsilateral projection represented respectively 49.2% and 43.5% of the cells (**Table 2**). There is a small contamination of the left spinal cord in the youngest E108 fetus (**Figure 2**), which however is not expected to influence the ipsilateral projecting CS neuron distribution. Nevertheless because of this contamination we rely principally on the older fetus for an account of the differences in the ipsi- and contralateral distributions of CS neurons (**Figure 5**).

**Table 2:** Number and percentage of neurons in the contralateral and ipsilateral CS pathway across ages.

| **Age** | **Hemisphere** | **Number of neurons** | **%** |
| --- | --- | --- | --- |
| E108 | Contralateral | 300324 | 50,8 |
| E108 | Ipsilateral | 290994 | 49,2 |
| E117 | Contralateral | 118422 | 56,5 |
| E117 | Ipsilateral | 91222 | 43,5 |
| Adult | Contralateral | 17496 | 79,9 |
| Adult | Ipsilateral | 4407 | 20,1 |

In the E117 fetus, CS neurons form a continuous band of back-labeled cells spanning the full extent of the frontal (green), parietal (black), insula (gray) and cingulate (yellow) regions. In the temporal cortex, CS neurons project from (i) the entire auditory subdivisions located within the posterior bank of the lateral sulcus (**Figure 5**, rows A to C), (ii) the presumptive temporal pole (**row B**), and (iii) the polysensory STP subdivisions within the anterior bank of the superior temporal sulcus. In the prefrontal cortex, ipsilateral CS neurons are present at lateral (**row A**), medial (**rows B** to **D**) and midline levels (row F), in presumptive areas 44, 45B, FEF and 8B/9. In the occipital cortex, no labeled CS neurons were found in area V1, and only very sparse labeling was observed in the vicinity of the occipital boundary (**row C**). In the limbic region, CS neurons were observed in the cingulate gyrus, in presumptive areas 23a/b, 24a/b, and 25/32 (not shown). Overall, at all ages and across ipsi- and contraleral hemispheres, the highest density of CS neurons was always found in the motor cortex, closely followed by the parietal cortex (**Figure 6**).

**Figure 6:**
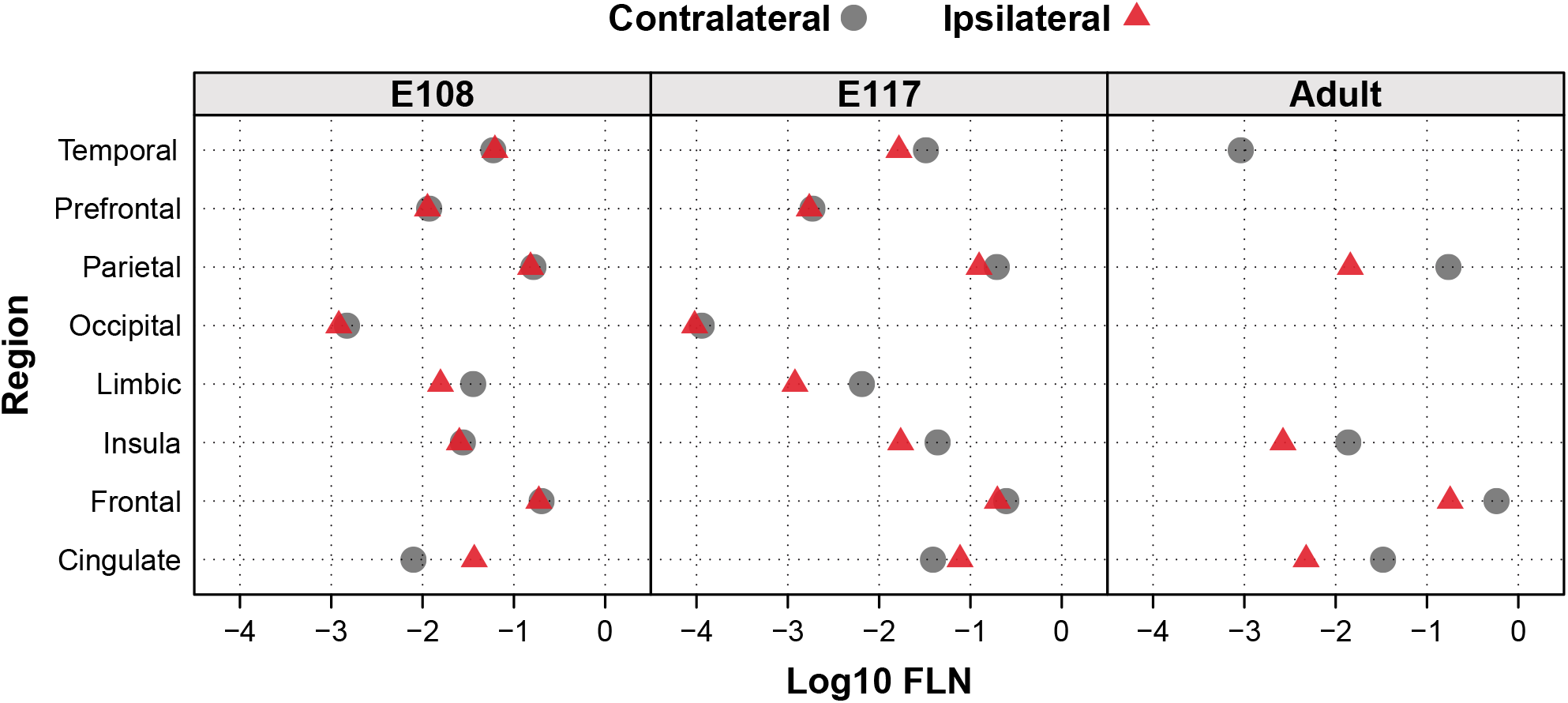
Relative strength of contralateral and ipsilateral projections across ages. Note the relative similarity of projections’ weights across all ages, the increasing differences in weights of contra- and ipsilateral projections with age and the complete loss of the prefrontal, occipital and limbic projections.

### Subcorticospinal pathways

In the present study we found abundant ipsi- and contralateral subcortical projections to the spinal cord with suggestive evidence of a developmental decrease in projections from the amygdala, hypothalamus and subthalamic nucleus (**Figure 7**). In the contralateral hemisphere, at all ages, the vast majority of back-labeled amygdospinal neurons projected from the central nucleus (**Figure 7, row A**). The projection from the hypothalamus (**row B**) was centered in the dorsal and posterior hypothalamic areas, but during development this projection was much broader, encompassing many hypothalamic subdivisions. The projection from the subthalamic nucleus to the cervical spinal cord (**row C**) was only observed in the fetuses, and no labeled neuron was observed in this nucleus in the adult. The distribution of the ipsilateral subcorticospinal projections is similar (not shown).

**Figure 7:**
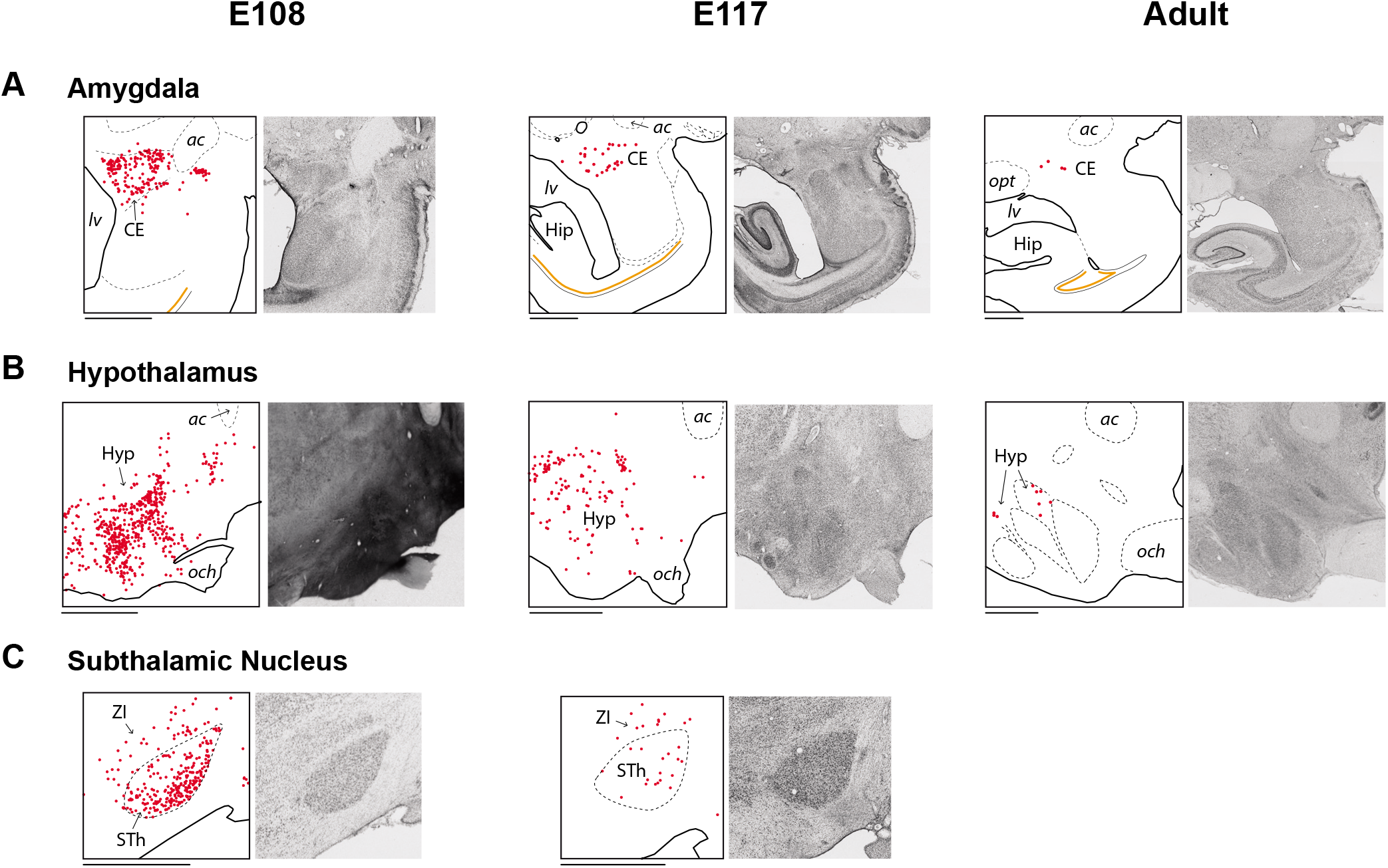
Subcorticospinal projections at E108, E117 and in the adult. Parasagittal sections of the left hemisphere showing the distribution of contralateral subcortical neurons projecting to the spinal cord from the amygdala (A), the hypothalamus (B) and the subthalamic nucleus (C). Abbreviations: ac, anterior commissure; CE, central nucleus of the amygdala; Hip, hippocampus; Hyp, hypothalamus; lv, lateral ventricle; och, optic chiasm; opt, optic tract; STh, subthalamic nucleus; ZI, zona incerta. Scale bar: 2 mm.

## Discussion

The progressive acquisition of manual dexterity in primates occurs in parallel with substantial changes in the underlying neuronal circuit indicating a link between functional and structural maturation during the first year of postnatal life (Kuypers 1962; Lawrence and Hopkins 1976; Galea and Darian-Smith 1995; Armand *et al.* 1997; Olivier *et al.* 1997). Despite the evolutionary importance of the CS pathway in primates, its early development has been substantially less investigated in comparison to non-primates. The six main findings of the present study, aimed at expanding our understanding of *in utero* primate CS pathway development are as follows. Firstly, in the prenatal macaque the somata of CS neurons in the contra- and ipsilateral hemispheres are restricted to layer 5. Secondly, the highest density of labeled CS neurons throughout development and in the adult is observed in the putative motor cortex. Thirdly, CS neurons are much more broadly distributed in the fetus, encompassing regions that are devoid of such neurons in the neonate and adult. Fourthly, while the adult CS pathway is strongly lateralized, during development there are comparable CS neurons in corresponding ipsi- and contralateral cortical territories. Fifthly, in parallel to the maturation of cortical projections to the spinal cord, there is substantial subcorticospinal maturation, whereby many projections to the spinal cord are eliminated from the amygdala, hypothalamus and subthalamic nuclei. Sixthly, the primary locus of the pick-up zone in the E117 fetus indicates an unexpected early and diffuse invasion of the gray matter of the spinal cord.

### Technical considerations

We have to rely on the visual identification of the pick-up zone, to infer the effective location of the injection site. Earlier investigation of the pick-up zone were carried out in the cortex, however we believe that similar results would be obtained for the spinal cord (Kennedy and Bullier 1985; Condé 1987). An indication that this is the case is the fact that following an injection up to the midline and which showed only a minute encroachment in the contralateral spinal cord, intense longitudinal labeling both anterior and posterior to the injection site was observed in the dorsal funiculus; importantly this longitudinal labeling was entirely restricted to the side of the spinal cord containing the pick-up zone (**Figure 1 and S1**). This argues in favor of the FB pick-up zone being largely restricted to the immediate vicinity of the needle tract as observed for cortical injections (Kennedy and Bullier 1985; Condé 1987).

The injection in the adult was carried out as in the fetus and intended to be restricted to one side of the spinal cord. In both fetuses and adult, injections were made under direct visual control using a high-power binocular surgical microscope. The surgery is considerably easier in the adult macaque and the quality of the visual control superior to that of the fetus. In the fetuses both injections were overall located as intended, the small pick-up zone in the contralateral cord in the E108 fetus likely caused by an error in handling of the micropipette subsequent to the correct injection.

In the adult the injection site material was not available for histological examination and so we have to rely on the labeling pattern to infer the location of the injection. There are significant differences in the topography of CS neurons reported in the primary motor and premotor areas following injections to the upper and lower cord levels (Liu and Chambers 1964; Kuypers and Brinkman 1970; He et al. 1993; Galea and Darian-Smith 1994, 1995). When the pick-up zone of the retrograde tracer includes the dorsolateral column, the cortical territory projecting to the injection site is expanded and includes regions that project not only to the level where the injection is located, but also lower levels of the spinal cord. In our adult material, cortical territories known to project to segments of the spinal cord posterior to the intended injected site were seen to contain many FB back-labeled neurons (**Figure 4, rows E and F**). Comparison with Galea and Darian-Smith (1994, 1995) suggests that the involvement of the dorsolateral funiculus is extensive. Further, reduced number of corticospinal neurons in the ipsilateral hemisphere is a strong indication that the injection was restricted to one side of the spinal cord, agreeing with the reduced ipsilateral CS projection described previously in the literature (Liu and Chambers 1964; Galea and Darian-Smith 1994, 1995; Dum and Strick 1996; Lacroix et al. 2004).

### There is a massive CS projection in the immature brain

The results from the present study show a massive cortical projection in the immature macaque brain to the cervical cord. This finding is in agreement with previous studies in non-primates. In rodents (rat: Leong (1983); Bates and Killackey (1984); Schreyer and Jones (1988)), carnivores (ferret: Meissirel *et al.* (1993)) and marsupials (opossum: Cabana and Martin (1984)), injections of retrograde tracers in the CS tract lead to the labeling of CS neurons distributed over a much broader cortical territory during development than in the mature animal, including transient projections from the prefrontal and occipital cortex.

Additionally, the results in the oldest fetus of the present study are of particular interest. Here, widespread projections were found following a discrete injection in the gray matter without encroaching on the dorsolateral funiculus and the ventral funiculus, where the descending CS tracts are housed. The widespread distribution of labeled CS neurons in this fetus suggests the existence of extensive transient axonal branching in the gray matter during development. This finding would seem to be at odds with the notion that there is a topological organization of the CS projection and that there is a progressive innervation of the spinal gray matter (Donatelle 1977). These findings reviewed by Stanfield (1992) indicate that in the rodent model there is a controversy concerning CS axons transiently entering into the spinal gray matter in regions where they are not present in the adult. A number of earlier studies in rodents report failure to label CS axons entering the spinal gray matter from cortical regions that do not have CS neurons in the adult (transient corticospinal axons) (Joosten and van Eden 1989; Joosten and Bar 1999) while others have found instances of innervation of spinal gray matter by transient CS axons (Stanfield and O’Leary 1985; O’Leary and Stanfield 1986). A recent retrograde study in mouse shows widespread projections to the gray matter of the cervical level of the spinal cord during CS pathway development (Kamiyama *et al.* 2015).

There are two series of observations that suggest that transient CS axons in primates may indeed enter into the spinal gray matter. Firstly, during development there is the possibility that individual CS fibers branch at multiple spinal levels, so that individual CS neurons in the adult could extend their axons to multiple spinal segments (Shinoda et al. 1981; He *et al.* 1993). Evidence of this occurs in cat where extensive spinal axonal collateralization with pruning of connections in latter stages is a feature of developing CS axons (Theriault and Tatton 1989). Secondly, there is some evidence of extensive innervation of the spinal gray matter observed in the postnatal monkey shortly after birth (Kuypers 1962; Armand *et al.* 1997). Likewise, innervation of the spinal gray matter by corticospinal projecting neurons has been suggested in postmortem studies in preterm human (Eyre et al. 2000).

Whether individual CS neurons innervate the spinal gray matter at multiple levels of the cord is extremely important for understanding the developmental refinement of the CS pathway. One could speculate that developmental pruning in the gray matter could lead to greater flexibility of the developmental outcome. Transient projections could be involved in activity dependent maturation of the spinal motor centers during a critical period of prenatal development, and the disruption of this process could lead to CS pathway perturbations that are observed following cortical lesions, such as those in cerebral palsy (Eyre *et al.* 2000; Eyre 2007). Note, the notion that transient projections fail to penetrate the gray matter has been argued in the cortex (Innocenti 1981); however, subsequently there has been irrefutable evidence in that such gray matter connections clearly exist (Dehay et al. 1984; Dehay, Kennedy and Bullier 1988). While the present results based on the location of the injection site in the E117 fetus and the apparent non-involvement of the dorsolateral funiculus are very indirect on this issue of innervation of the cord, they point to the need for further studies on the development of the CS pathway using modern anterograde tracer injection in diverse regions of the cortex in areas that do and do not maintain permanent CS projections in the adult. Exploring the extent of prenatal gray matter innervation in primates will address the extent to which the specificity of the developmental refinement of the primate CS pathway resides in the details of its timing, and/or in the pattern of gray matter innervation.

### Is there a homogeneous distribution of CS neurons in the prenatal brain?

To address this issue we need to allocate labeled neurons to particular areas or regions. At present there is no available atlas for the developing macaque cortex, and only a handful of publications have investigated macaque area identification during fetal development (Goldman and Galkin 1978; Dehay, Kennedy, Bullier, et al. 1988; Dehay et al. 1989; Dehay et al. 1993; Barone et al. 1994; Dehay et al. 1996). This led us to combine cytoarchitectonics with sulci and gyri landmarks in the E108 and E117 to define in the developing cortex, 8 different cortical territories which can be readily identified in the adult cortex (prefrontal, frontal, insular, temporal, parietal, occipital, cingulate and limbic). Within these regions, as early as E108 the boundaries of the primary visual and auditory areas can be detected.

Having identified these cortical regions across all fetal and adult stages we are able to determine the developmental decrease in CS neuron number. This showed that the pattern of labeling across regions remains remarkably stable (**Figure 6**). Overall, during development and in the adult macaque, the strongest CS projection arises from the frontal cortex and the second strongest from the parietal cortex, followed by the cingulate and insular cortices (**Figure 6**). This demonstrates that prenatally, the regions projecting to the spinal cord are already well defined in terms of differences in numbers of CS neurons.

### Exuberant bilateral corticospinal projections

In the present study we show that in primates, as for the rodent, carnivores, and marsupials, there is a developmental narrowing of the extent of the cortex projecting to the spinal cord (Stanfield *et al.* 1982; Leong 1983; Bates and Killackey 1984; Cabana and Martin 1984; O’Leary and Stanfield 1986; Sharkey et al. 1986; Joosten et al. 1987; Meissirel *et al.* 1993). This is not only true for the regions that project in the adult and neonate (He *et al.* 1993; Galea and Darian-Smith 1994, 1995; He *et al.* 1995), but also from regions that do not have CS neurons postnatally. These include the prefrontal, occipital, limbic and temporal cortices. This exuberance was found bilaterally. While in the adult macaque there is a dominant contralateral CS projection, this does not appear to be the case prenatally.

During development of the CS pathway in the macaque we observed an extensive ipsilateral projection to the cervical cord. Further, the distribution pattern of back-labeled neurons is comparable across the ipsi- and contralateral hemispheres (**Figures 4 and 5 and Table 2**). Most studies investigating CS pathway development in non-primates employed anterograde tracer injections in the cortex and investigated the spatial distribution of the terminal labeling at different spinal levels. Studies in cat show that there are extensive bilateral terminations of CS axons with fairly similar terminal distribution patterns (Alisky *et al.* 1992; Li and Martin 2000). The Li and Martin (2000) study showed that although the density of labeling in the ipsi- and contralateral gray matter exhibits an important degree of variability, with the ipsilateral label densities ranging from 11 to 50% of the total amount of labeling on a section of the cord, the topography of ipsilateral labeling is more consistent. Hence, unilateral injections of retrograde tracer in kittens could produce relatively similar patterns of labeled neurons in the ipsi- and contralateral hemispheres, in agreement with the findings of the present study.

### Exuberant bilateral subcorticospinal projections

A number of studies have examined non-cortical origins of projections to the spinal cord in the adult macaque by (1) lesioning descending fiber pathways, (2) injecting anterograde tracers in different subcortical structures, and (3) injecting retrograde tracers in the spinal cord (e.g. Kneisley et al. 1978). These studies mainly focused on the projections from the midbrain and hindbrain (reviewed in Lemon 2008), namely those from the magnocellular red nucleus (rubrospinal), the superior culliculus (tectospinal), the vestibular nuclei (vestibulospinal) and the pontine and medullary reticular formation (reticulospinal).

Additional projections were found from other subcortical structures in the adult brain, but subject to a much less intensive investigation. Very weak, uniquely ipsilateral amygdalospinal projections from the central nucleus of amygdala were described by Mizuno et al. (1985), similarly weak and ipsilateral projections from the hypothalamus were described by Kneisley *et al.* (1978) and Barbas et al. (2003) and a few subcorticospinal projecting neurons from the subthalamic nucleus were shown by Mizuno et al. (1988).

In the present study we observed subcorticospinal projecting neurons bilaterally in the central amygdala and in the hypothalamus of the adult and during development, and we found in the fetues that these projections appeared considerably more extensive then in the adult (**Figure 7**). In addition, we observed a transient projection from the subthalamic nucleus to the spinal cord; no labeled neuron was found in our adult case. To our knowledge these pathways were not described during development in other species. Future studies would be needed to determine if these pathways are present in non-primates, and if they are reorganized following early lesions, as observed for CS projections.

### Prenatal lesions and aberrant ipsilateral CS projections

Experimental manipulations of the CS system in rodent suggest an important degree of plasticity of the projections from the cortex to the spinal cord. These findings are potentially important for clinical observations. First, occipital neurons that are transplanted at early stages of development to the frontal cortex, retain their transient projection to the spinal cord (O’Leary and Stanfield 1985). Second, lesions of the sensorimotor cortex during early development lead to an increase in the ipsilateral projections from the non-lesioned cortex, with similar pattern of the contralateral projection (Reinoso and Castro 1989). While these findings have not been replicated in primate, lesion studies in perinatal human point to similar processes. Prenatal lesions in humans lead to hypertrophy of the CS tract arising from the undamaged (non-affected) hemisphere, which projects to both sides of the spinal cord and innervates motoneuronal pools (Eyre 2007).

The results of our study demonstrate that during primate development there is a bilateral projection from both ipsi- and contralateral hemispheres to the spinal cord and further that both sets of projections have relatively similar strengths. It could be speculated that the increased ipsilateral projection in human (Eyre 2007), results from the maintenance of the transient ipsilateral projection. A lesion to the cortex may lead to impairments of the normal activity and motor experience-dependent mechanisms of axonal pruning, with subsequence imbalance of ipsilateral pruning (Martin 2005; Eyre 2007).

## Conclusion

The importance of the macaque model for understanding health issues with respect to spinal cord injury is well established (Courtine et al. 2007). Further, on top of species-specific motor system organization, comparisons across species is problematic because retrograde and anterograde studies can give very different results according to the location of injection site, the tracer used and the developmental stage examined. The recent description of a transient projection of the cortex to motoneuron pools in the rodent (Maeda et al. 2016; Gu et al. 2017; Murabe et al. 2018), cautions against simplification of primate specific features. This is of particular relevance to clinical studies in human which address the medical outcome of disturbed development of the CS pathway (Eyre 2007; Welniarz *et al.* 2017). These considerations point to the need for an in-depth study of the development of the primate CS pathway.

## Acknowledgements

Funded by LABEX CORTEX (ANR-11-LABX-0042) of Université de Lyon (ANR-11-IDEX-0007) operated by the French National Research Agency (ANR) (H.K.), ANR-15-CE32-0016, CORNET (H.K.), ANR-17-NEUC-0004, A2P2MC (H.K.), ANR-17-HBPR-0003, CORTICITY (H.K.), Chinese Academy of Sciences President’s International Fellowship Initiative. Grant No. 2018VBA0011 (H.K.), FRC “Rotary-Espoir en Tête” 2018 (H.K.).

## Author Contributions

Surgery PG, MB, CD, HK; Histological processing ARRG, PG, CD; Data acquisition ARRG, EO, HK; Data analysis and discussion/interpretation ARRG, EO, KK, CD, HK; Wrote the paper ARRG, EO, KK, CD, HK; All authors revised successive versions of the manuscript.

