## Supplementary Figure S1 and Table S1 for "Refinement of the primate corticospinal pathway during prenatal development"

### Supplementary figure legend

**Figure S1: E117 Injection site**. Horizontal sections of the upper spinal cord and medulla with the pick-up zone of the injection site indicated in red, accompanied by high-exposure micrographs caudal to the injection site highlighting the labeled fibers limited to the right hand side of the spinal cord. Black and white arrows highlight the blood vessels present along the midline, across the ventral to dorsal aspects of the spinal cord. Scale bar: 2 mm.

### Supplementary table legend

**Table S1: Allocation of areas to regions**.

### Figure S1

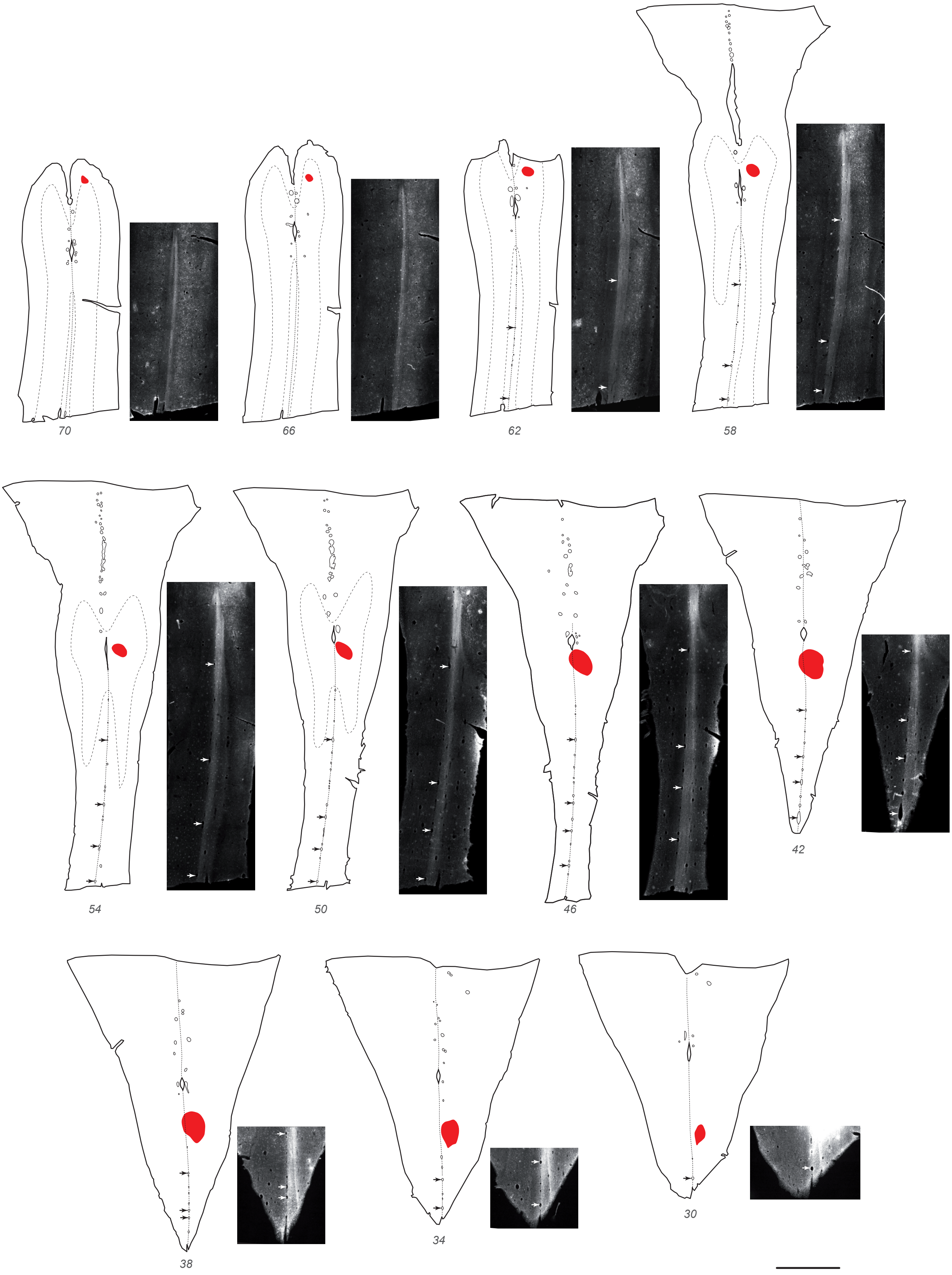

*Table S1: Allocation of areas to regions*

| Abbrev. | Area name | Region |
|---|---|---|
| TEO | Temporal area TE, occipital part (occipito-temporal juntion) | Occipital |
| V1 | Visual area 1 | Occipital |
| V2 | Visual area 2 | Occipital |
| V3 | Visual area 3 | Occipital |
| V3A | Visual area 3, part A | Occipital |
| V4 | Visual area 4 | Occipital |
| Aud. Core | Auditory core (includes the primary auditory cortex) | Temporal |
| Entorhinal | Entorhinal | Temporal |
| FST | Fundus of the superior temporal sulcus | Temporal |
| IPa | Intraparietal sulcus associated area in the superior temporal sulcus | Temporal |
| LB | Belt region of the auditory cortex, lateral part | Temporal |
| MB | Belt region of the auditory cortex, medial part | Temporal |
| MST | Medial superior temporal area | Temporal |
| MT | Middle temporal area | Temporal |
| PBc | Parabelt region of the auditory cortex, caudal part | Temporal |
| PBr | Parabelt region of the auditory cortex, rostral part | Temporal |
| Perirhinal | Perirhinal | Temporal |
| PGa | PG associated area of the superior temporal sulcus | Temporal |
| Piriform | Piriform | Temporal |
| Pro.St. | Prostriata | Temporal |
| STPc | Superior temporal polysensory area, caudal subdivision | Temporal |
| STPi | Superior temporal polysensory area, intermediary subdivision | Temporal |
| STPr | Superior temporal polysensory area, rostral subdivision | Temporal |
| Subic.Comp. | Subicular complex (Parasubiculum, presubiculum, prosubiculum and subiculum) | Temporal |
| TE | Temporal area TE (TEm/a, TEad, TEav, TEpd, TEpv) | Temporal |
| T.Pole | Temporal pole | Temporal |
| TEOm | Temporal area TE, occipitomedial part | Temporal |
| TH/TF | Areas TH and TF of the parahippocampal cortex | Temporal |
| TPt | Temporoparietal area (temporoparietal junction) | Temporal |
| V4t | Visual area 4, transitional part | Temporal |
| 1 | Somatosensory area 1 | Parietal |
| 2 | Somatosensory area 2 | Parietal |
| 3a | Somatosensory area 3a | Parietal |
| 3b | Somatosensory area 3b, primary somatosensory cortex | Parietal |

| Abbrev. | Area name | Region |
|---|---|---|
| 5 | Somatosensory area 5 | Parietal |
| 7A | Area 7A (caudal inferior parietal lobule area) | Parietal |
| 7B | Area 7B (rostral inferior parietal lobule area) | Parietal |
| 7m | Area 7m (medial parietal area) | Parietal |
| 7op | Area 7op (parietal operculum) | Parietal |
| AIP | Anterior intraparietal area | Parietal |
| DP | Dorsal prelunate area | Parietal |
| LIP | Lateral intraparietal area | Parietal |
| MIP | Medial intraparietal area | Parietal |
| PIP | Posterior intaparietal area | Parietal |
| V6 | Visual area 6 (parieto-occipital area) | Parietal |
| V6A | Visual area 6A (parieto-occipital area) | Parietal |
| VIP | Ventral intraparietal sulcal area | Parietal |
| SII | Secondary somatosensory area (parietal operculum) | Parietal |
| Insula | Insula | Insula |
| F1 | Agranular frontal area 1, primary motor cortex | Frontal |
| F2 | Agranular frontal area 2 (dorsal posterior premotor area) | Frontal |
| F3 | Agranular frontal area 3 (supplementary motor area) | Frontal |
| F4 | Agranular frontal area 4 (ventral posterior premotor area) | Frontal |
| F5 | Agranular frontal area 5 (ventral anterior premotor area) | Frontal |
| F6 | Agranular frontal area 6 (pre-supplementary motor area) | Frontal |
| F7 | Agranular frontal area 7 (dorsal anterior premotor area) | Frontal |
| Gu | Gustatory cortex (frontal operculum) | Frontal |
| ProM | Area ProM (promotor) (frontal operculum) | Frontal |
| 10 | Area 10 (frontal pole) | Prefrontal |
| 11 | Area 11 (orbitofrontal cortex) | Prefrontal |
| 12 | Area 12 | Prefrontal |
| 13 | Area 13 (orbitofrontal cortex) | Prefrontal |
| 14 | Area 14 | Prefrontal |
| 44 | Area 44 | Prefrontal |
| 45A | Area 45A | Prefrontal |
| 45B | Area 45B | Prefrontal |
| 46d | Area 46, dorsal part (dorsolateral prefrontal cortex) | Prefrontal |
| 46v | Area 46, ventral part (dorsolateral prefrontal cortex) | Prefrontal |
| 8B | Area 8B | Prefrontal |
| 8r | Area 8, rostral part | Prefrontal |
| 9 | Area 9 | Prefrontal |
| 9/46d | Area 9/46, dorsal part (dorsolateral prefrontal cortex) | Prefrontal |
| 9/46v | Area 9/46, ventral part (dorsolateral prefrontal cortex) | Prefrontal |
| FEF | Frontal eye field (area 8, lateral part and medial part) | Prefrontal |

| **Abbrev.** | **Area name** | **Region** |
|---|---|---|
| 23c | Area 23, subdivisions c | Cingulate |
| 24c | Area 24, subdivisions c | Cingulate |
| 24d | Area 24, subdivisions d | Cingulate |
| 23a/b | Area 23, subdivisions a and b | Limbic |
| 24a/b | Area 24, subdivisions a and b | Limbic |
| 25 | Area 25 | Limbic |
| 29/30 | Areas 29 and 30 of the retrosplenial cortex | Limbic |
| 31 | Area 31 | Limbic |
| 32 | Area 32 | Limbic |
